# Population structure and coalescence in pedigrees: comparisons to the structured coalescent and a framework for inference

**DOI:** 10.1101/054957

**Authors:** Peter R. Wilton, Pierre Baduel, Matthieu M. Landon, John Wakeley

**Author notes:** Corresponding author: *Email address:* (Peter R. Wilton).

## Abstract

Contrary to what is often assumed in population genetics, independently segregating loci do not have completely independent ancestries, since all loci are inherited through a single, shared population pedigree. Previous work has shown that the non-independence between gene genealogies of independently segregating loci created by the population pedigree is weak in panmictic populations, and predictions made from standard coalescent theory are accurate for populations that are at least moderately sized. Here, we investigate patterns of coalescence in pedigrees of structured populations. We find that the pedigree creates deviations away from the predictions of the structured coalescent that persist on a longer timescale than in the case of panmictic populations. Nevertheless, we find that the structured coalescent provides a reasonable approximation for the coalescent process in structured population pedigrees so long as migration events are moderately frequent and there are no migration events in the recent pedigree of the sample. When there are migration events in the recent sample pedigree, we find that distributions of coalescence in the sample can be modeled as a mixture of distributions from different initial sample configurations. We use this observation to motivate a maximum-likelihood approach for inferring migration rates and mutation rates jointly with features of the pedigree such as recent migrant ancestry and recent relatedness. Using simulation, we show that our inference framework accurately recovers long-term migration rates in the presence of recent migration events in the sample pedigree.

## 1. Introduction

The coalescent is a stochastic process that describes the distribution of gene genealogies, the tree-like structures that describe relationships among the sampled copies of a gene. Since its introduction (Kingman, 1982a,b; Hudson, 1983; Tajima, 1983), the coalescent has been extended and applied to numerous contexts in population genetics and is now one of the foremost mathematical tools for modeling genetic variation in samples (Hein *et al.*, 2005; Wakeley, 2009).

In a typical application to data from diploid sexual organisms, the coalescent is applied to multiple loci that are assumed to have entirely independent ancestries because they are found on different chromosomes and thus segregate independently, or are far enough apart along a single chromosome that they effectively segregate independently. Even chromosome-scale coalescent-based inference methods that account for linkage and recombination (e.g., Li and Durbin, 2011; Sheehan *et al.*, 2013; Schiffels and Durbin, 2014) multiply probabilities across distinct chromosomes that are assumed to have completely independent histories due to their independent segregation.

In reality, the ancestries of gene copies sampled at independently segregating loci from a fixed set of diploid, sexually reproducing individuals are independent only after conditioning on the population pedigree, i.e., the set of familial relationships between all individuals in the population throughout all time. Figure 1 illustrates this. Figure 1A depicts coalescence within the framework of standard coalescent theory under a diploid, monoecious Wright-Fisher model with the possibility of selfing. Under this model, from which the coalescent can be derived as a limiting process, the probability that two distinct ancestral lineages coalesce in a given generation is 1*/*(2*N*), where *N* is the diploid population size. The averaging over pedigrees in this model is depicted in Figure 1A by each individual having a “parental” relationship to every individual in the previous generation. Thus, two ancestral lineages follow parental relationships to the same individual with probability 1*/N*, and in that individual are derived from the same parental chromosome with probability 1*/*2. When this probability, 1*/*(2*N*), is applied independently to unlinked loci, any correlation in ancestry between these loci caused by the pedigree is erased by this averaging over pedigrees.

**Figure 1:**
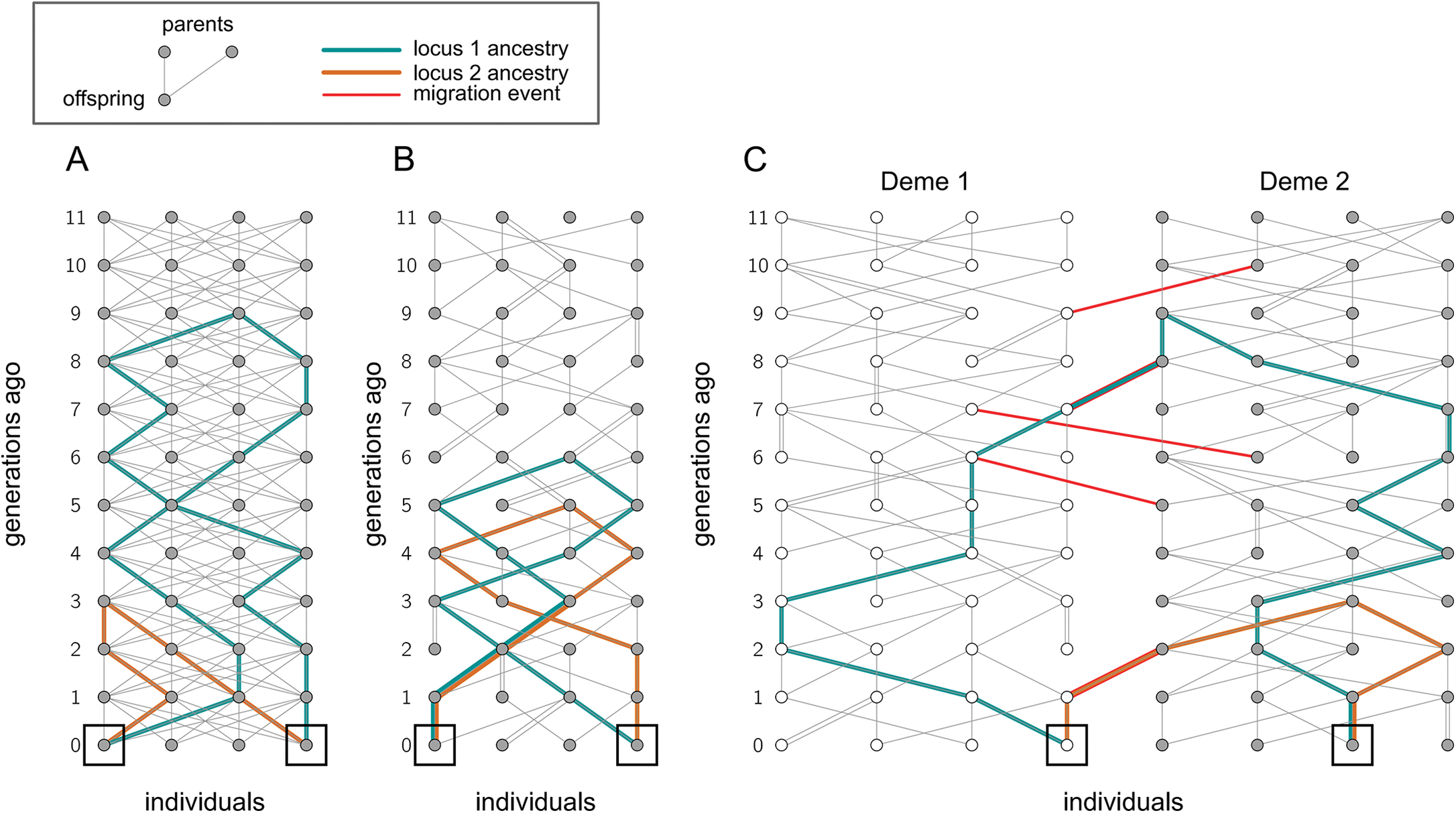
A conceptual inconsistency in the application of standard coalescent theory to independently segregating loci sampled from a diploid population. Each panel depicts coalescence between two sampled gene copies at a pair of unlinked loci (green and orange lines, respectively) sampled from a pair of individuals in the present generation (black squares). Individuals within a generation are represented by circles arranged horizontally across a row, and generations are arranged vertically, with the present generation at the bottom of the figure and the most ancient generation at the top of the figure. Panel **A)** is a conceptual depiction of coalescence at unlinked loci under the assumptions of standard coalescent theory. Under these assumptions, at each unlinked locus, an ancestral lineage has an equal probability of being derived from any individual in the previous generation, and thus it is as if each individual has every potential ancestral individual as a parent. Panels **B)** and **C)** depict coalescence at two unlinked loci in actual diploid sexual populations, in which the same population pedigree governs the process of coalescence at all loci. In these pedigrees, each individual has exactly two parents (including the possibility of selfing, here), and the distribution of coalescence times amongst unlinked loci depends on both the initially sampled individuals and the particular shape of the pedigree. Panel **B)** shows the pedigree of a panmictic population. The probability of coalescence of two ancestral lineages in a given generation depends on the pedigree and generally differs from the 1*/*(2*N*) that would be assumed after marginalizing over the pedigree. Panel **C)** depicts a two-deme population pedigree, with fixed migration events (red lines) constraining the movement of ancestral lineages between the two demes.

A conceptually more correct model of coalescence of unlinked loci sampled from individuals in a diploid sexual population is depicted in Fig. 1B. Here, coalescence occurs within a single diploid population pedigree, with probabilities of coalescence in a given generation determined by the particular shape of the pedigree of the sampled individuals and generally differing from the 1*/*(2*N*) predicted under the standard interpretation of the diploid Wright-Fisher model. For example, if in a given generation the number of shared parental relationships happens to be greater than usual, the probability of coalescence will tend to be greater, and when the number of shared parental relationships in a given generation is fewer, the probability of coalescence in that generation will tend to be lesser.

These effects of the population pedigree on coalescence in sexual populations were investigated by Wakeley *et al.* (2012), who studied coalescence in randomly generated population pedigrees of diploid populations reproducing under basic Wright-Fisher-like dynamics, *i.e.*, populations with constant population size, random mating, non-overlapping generations, and no population structure, and in this context, it was found that the shape of the population pedigree affects coalescence probabilities mostly during the first ∼ log_2_(*N*) generations back in time. During these first few generations back in time, there is relatively large variation from pedigree to pedigree in the degree of overlap in the pedigree of the sampled individuals. It may be that for several generations there is zero overlap in the pedigree of the sampled individuals, and consequently the probability of coalescence amongst these individuals will be zero. On the other hand, if by chance the sample contains close relatives (i.e., two individuals possessing a shared pedigree ancestor in these recent generations), the probability of coalescence in the generation of that shared ancestor will tend to exceed the probability of coalescence predicted by standard theory. After the first log_2_(*N*) generations, the pedigrees of different individuals begin to overlap more completely, and the probabilities of coalescence in these later generations depend less on the pedigree (Wakeley *et al.*, 2012, see also Fig. 2).

This log_2_(*N*) timescale of convergence or mixing of the pedigree has been studied in other contexts. Chang (1999) found that the number of generations until two individuals share an ancestor (in the biparental pedigree sense) converges to log_2_(*N*) as the population size grows. Likewise, Derrida *et al.* (2000) showed that the distribution of the number of repetitions in an individual’s pedigree ancestry becomes stationary around log_2_(*N*) generations in the past. This log_2_(*N*)-generation timescale is the natural timescale of convergence in pedigrees due to the approximate doubling of the number of possible ancestors each generation back in time until the entire ancestral population potentially becomes part of the pedigree.

In these studies it is assumed that the population is panmictic, *i.e.*, that individuals mate with each other uniformly at random. One phenomenon that may alter this convergence in pedigrees is population structure, with migration between subpopulations or demes. In a subdivided population, the exchange of ancestry between demes depends on the history of migration events embedded in the population pedigree (Fig. 1C). These fixed past migration events may be infrequent or irregular enough that the generation-by-generation probabilities of coalescence depend on the details of the migration history rather than on the reproductive dynamics underlying convergence in panmictic populations.

The question of how population structured affects pedigree ancestry has received some previous attention. Rohde *et al.* (2004) found that population structure did not change the log_2_(*N*)-scaling of the number of generations until a common ancestor of everyone in the population is reached. Barton and Etheridge (2011) studied the expected number of descendants of an ancestral individual, a quantity termed the reproductive value, and similarly found that population subdivision did not much slow the convergence of this quantity over the course of generations. Kelleher *et al.* (2016) studied how pedigree ancestry spreads in spatially explicit models of population structure, finding that when dispersal is local, the mixing of pedigree ancestry occurs on a longer timescale than the log_2_(*N*)-scaling found by Rohde *et al.* (2004).

While these studies give a general characterization of how pedigrees are affected by population structure, a direct examination of the coalescent process for loci segregating through the pedigree of a structured population is still needed. Fixing the migration events in the pedigree may produce long-term fluctuations in coalescence probabilities that make the predictions of the structured coalescent inaccurate. The pedigree may also bias inference of demographic history. Any migration events or overlap in ancestry in the most recent generations of the sample pedigree will have a relatively great probability of affecting sampled genetic variation, and thus demographic inference methods that do not take into account the pedigree of the sample may be biased by how these events shape genetic variation in the sample.

Here, we explore how population structure affects coalescence through randomly generated population pedigrees. Using simulations, we investigate how variation in the migration history embedded in the pedigree affects coalescence probabilities, and we determine how these pedigree effects depend on population size and migration rate. We also study the effects of recent migrant ancestry on patterns of coalescence and use our findings to develop a simple framework for modeling the sample as a probabilistic mixture of multiple ancestries without migration ancestry. We demonstrate how this framework can be incorporated into demographic inference by developing a maximum-likelihood method of inferring scaled mutation and migration rates jointly with the recent pedigree of the sample in a population with two demes and migration between the demes. We test this inference approach with simulations, showing that including the pedigree in inference corrects a bias that is present when there is unaccounted-for migration in the ancestry of the sample.

## 2. Theory and Results

### 2.1. Pedigree simulation

Except where otherwise noted, each population we consider has two demes of constant size, exchanging migrants symmetrically at a constant rate. This model demonstrates the effects of population structure in one of the simplest ways possible. Under the assumptions of standard coalescent theory, coalescence in such a two-deme population is described by the structured coalescent (Notohara, 1990; Hudson, 1991), a model describing migration of ancestral lineages between demes and coalescence of ancestral lineages within a deme. With just two demes and a constant rate of migration, this model has a relatively simple mathematical description (Wakeley, 2009).

We assume that generations are non-overlapping and that the population has individuals of two sexes in equal number. In each generation, each individual chooses a mother uniformly at random from the females of the same deme with probability 1 − *m* and from the females of the other deme with probability *m*. Likewise a father is chosen uniformly at random from the males of the same deme with probability 1 − *m* and from the males of the other deme with probability *m*. This system of reproduction corresponds to reproduction by broadcast spawning, where gametes migrate but individuals do not. This migration scheme maintains a constant deme size from generation to generation.

In our depictions of pedigrees in the figures of this paper, we do not distinguish between males and females, as none of our results depend on the sexes of the individuals. The present generation, from which individuals are always sampled, is depicted at the bottom of the pedigree, with the two lines extending upwards showing the maternal and paternal relationships to the previous generation. When there are two demes, individuals inhabiting one or the other deme are colored white or gray, respectively.

All simulations were carried out with `coalseam`, a program for simulation of coalescence through randomly generated population pedigrees. The user provides parameters such as population size, number of demes, mutation rate, and migration rate, and coalseam simulates a population pedigree under a Wright-Fisher-like model meeting the specified conditions. Gene genealogies are constructed by simulating segregation back in time through the pedigree, and the resulting genealogies are used to produce simulated genetic loci. Output is in a format similar to that of the program ms (Hudson, 2002), and various options allow the user to simulate and analyze pedigrees featuring, for example, recent selective sweeps or fixed amounts of recent pedigree relatedness and migrant ancestry.

The program `coalseam` is written in C and released under the GPLv3 license. It is available online at https://github.com/ammodramus/coalseam.

**Figure 2:**
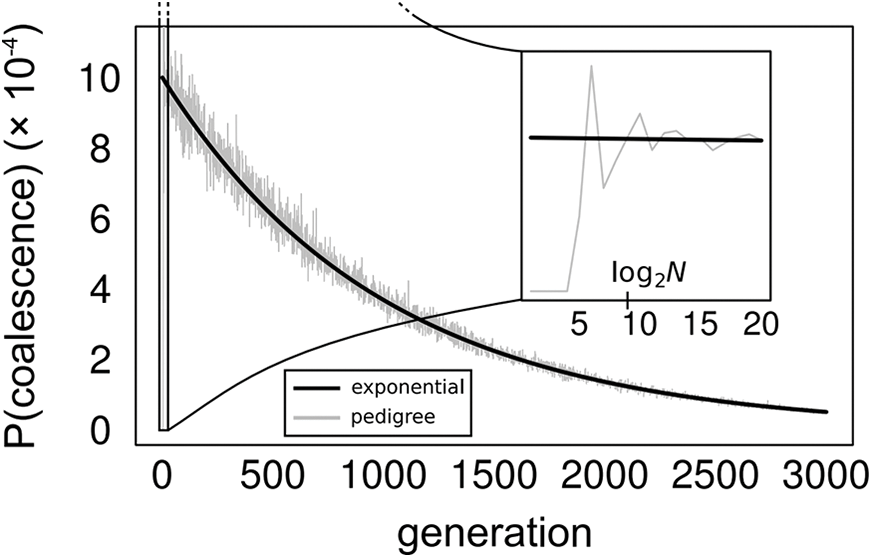
Coalescence time distribution for independently segregating loci sampled from two individuals in a panmictic population. The gray line shows the distribution from the pedigree, and the black line shows the exponential prediction of the standard coalescent. The population size is *N* = 500 diploid individuals.

### 2.2. Structured population pedigrees and probabilities of coalescence

As described by (Wakeley *et al.*, 2012), in a well-mixed population of size *N*, the distribution of coalescence times for independently segregating loci sampled from two individuals shows large fluctuations over the first ∼ log_2_(*N*) generations depending on the degree of overlap in the pedigree of the two individuals. After this initial period, the coalescence probabilities quickly converge to the expectation under standard coalescent theory, with small fluctuations around that expectation (Fig. 2, see also Wakeley *et al.* 2012). The magnitude of these fluctuations depends on the population size, but even for small to moderately sized populations (e.g., *N* = 500), the exponential prediction of the standard coalescent is a good approximation to the true distribution after the first log_2_(*N*) generations.

In a large structured population of two populations with symmetric migration rate *m* per generation and diploid deme size *N*, the coalescent process is described by the two-deme structured coalescent with scaled migration rate *M/*2 = 2*Nm* per lineage and coalescence rate 1 for every pair of lineages in the same deme (Wakeley, 2009). The distribution of pairwise coalescence times under this model can be calculated by exponentiating a matrix of migration and coalescence rates, giving the distribution of coalescence times for two chromosomes sampled within the same deme,

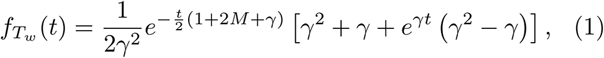

and the distribution of coalescence times for two chromosomes sampled from different demes,

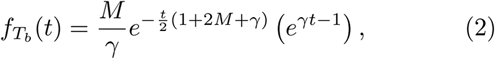

where *t* is *measured* in units of 2*N* generations and 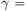
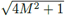. These are the predicted distributions of pairwise coalescence times in a two deme population using the standard approach that implicitly involves marginalizing over pedigree relationships.

When the pedigree of a two-deme population is fixed, coalescence probabilities depend on the history of migration embedded in the population pedigree. Deviations from the predictions of the structured coalescent are especially pronounced when the average number of migration events per generation is of the same order as the per-generation pairwise coalescence probability, i.e., *N m ≈* 1*/N* (Fig. 3A). In this migration-limited regime, two lineages may be “stuck” in different demes and have zero probability of coalescing before a migration event in the pedigree can bring them together into the same deme. This creates large peaks in the coalescence time distributions for loci segregating independently through the same pedigree, with each peak corresponding to a migration event (Fig. 3A).

**Figure 3:**
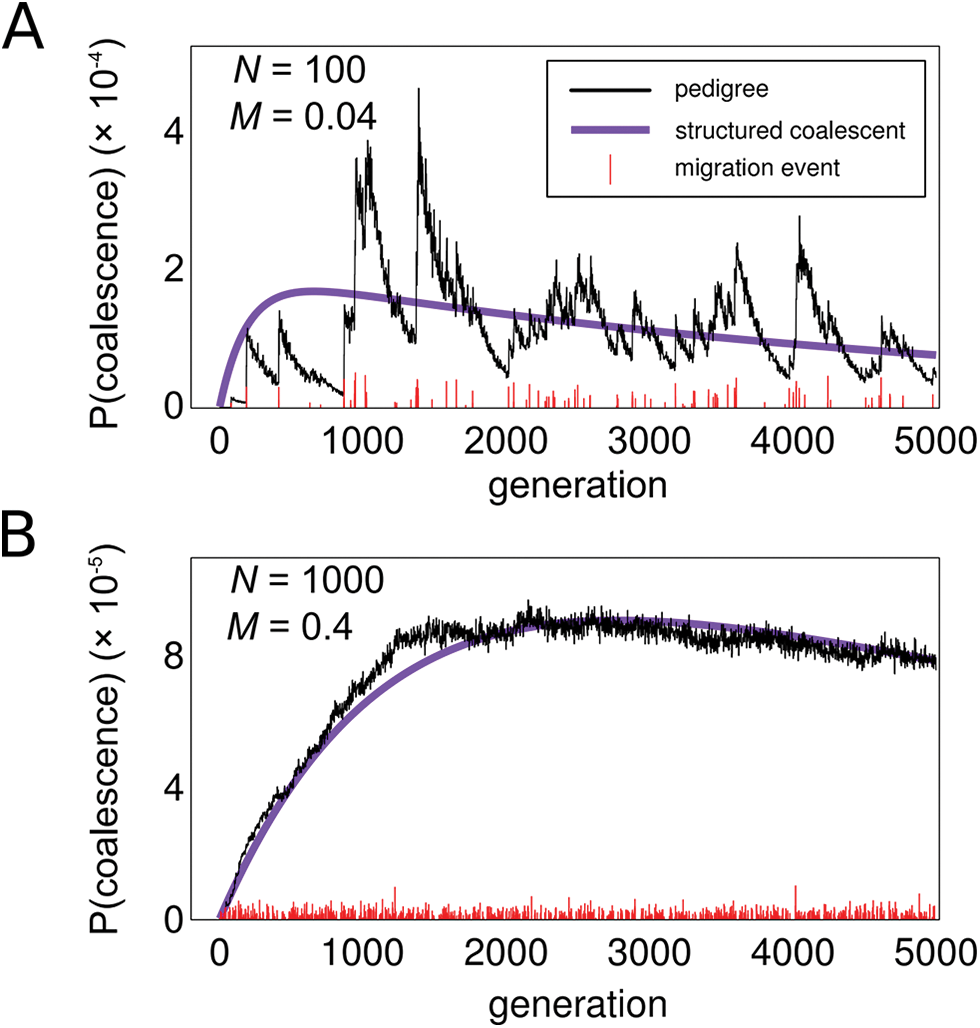
Distribution of coalescence times for two individuals sampled from different demes. In both panels, the black line shows the distribution calculated from a single random pedigree, and the purple line shows the prediction from the structured coalescent. Red vertical lines along the horizontal axis show the occurrence of migration events in the population, with the relative height representing the total reproductive weight (see Barton and Etheridge, 2011) of the migrant individual(s) in that generation. **A)** Low migration pedigree, with *M* = 4*Nm* = 0.04 and *N* = 100. Under these conditions, coalescence is limited by migration events, so there are distinct peaks in the coalescence time distribution corresponding to individual migration events. **B)** Higher migration pedigree, with *M* = 0.4 and *N* = 1000. With the greater migration rate, coalescence is no longer limited by migration, but stochastic fluctuations in the migration process over time cause deviations away from the standard-coalescent prediction on a timescale longer than log_2_(*N*) generations.

Even when the migration rate is greater and there are many migration events per coalescent event, the pedigree can still cause coalescence probabilities to differ from the predictions of the structured coalescent. Under these conditions, coalescence is not constrained by individual migration events, but there may be stochastic fluctuations in the realized migration rate, with some periods experiencing more migration and others less. These fluctuations can cause deviations in the predicted coalescence probabilities long past the log_2_(*N*)-generation timescale found in well-mixed populations (Figs. 3B, S1, S2). The degree of these deviations depends on the rate of migration and the population size, with smaller populations and lower migration rates causing greater deviations, and deviations from predictions are generally larger for samples taken between demes than samples within demes. Overall, however, when there are many migration events per coalescent event (i.e., when 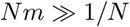), the predictions of the structured coalescent fit the observed distributions in pedigrees reasonably well (Figs. S1–S2).

To investigate the dependence of the coalescence time distribution on the pedigree more systematically, we simulated 20 replicate population pedigrees with different deme sizes and migration rates. From each pedigree, we sampled two individuals in different demes and calculated the distribution of pairwise coalescence times for independently segregating loci sampled from those two individuals. We measured the total variation distance of this distribution from the distribution of coalescence times predicted under a two-deme discrete-time Wright-Fisher model with migration rate equal to that of the simulation. The total variation distance of two discrete distributions *P* and *Q* is defined as

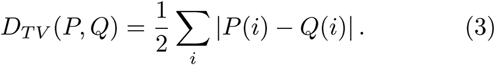

In theory this is an infinite sum, but in practice, we sum through 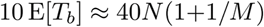 generations (where 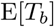 is the expected between-deme coalescence time), beyond
which there is very little probability remaining in these distributions and thus very little additional contribution to the total variation distance. We find that the total variation distance between the distributions from pedigrees and the distributions from standard theory increases as both the population size and migration rate decrease (Fig. 4A). When the total, population-scaled rate of migration (i.e., *Nm*) is greater than the rate of coalescence within demes (i.e., when 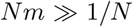, or 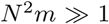, dotted line in Fig. 4B), total variation distance depends more on the population size than on the migration rate. When the total migration rate is less than the rate of coalescence within demes (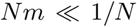), total variation distance increases greatly and depends more on the population-wide migration rate than on the deme size (Fig. 4B). This shows that the pedigree has strong effects on coalescence when the average number of migrants per generation, *Nm*, is relatively small compared to the per-generation probability of coalescence within a deme, 1*/*(2*N*). In this case, patterns of coalescence are not well approximated by typical coalescent theory.

**Figure 4:**
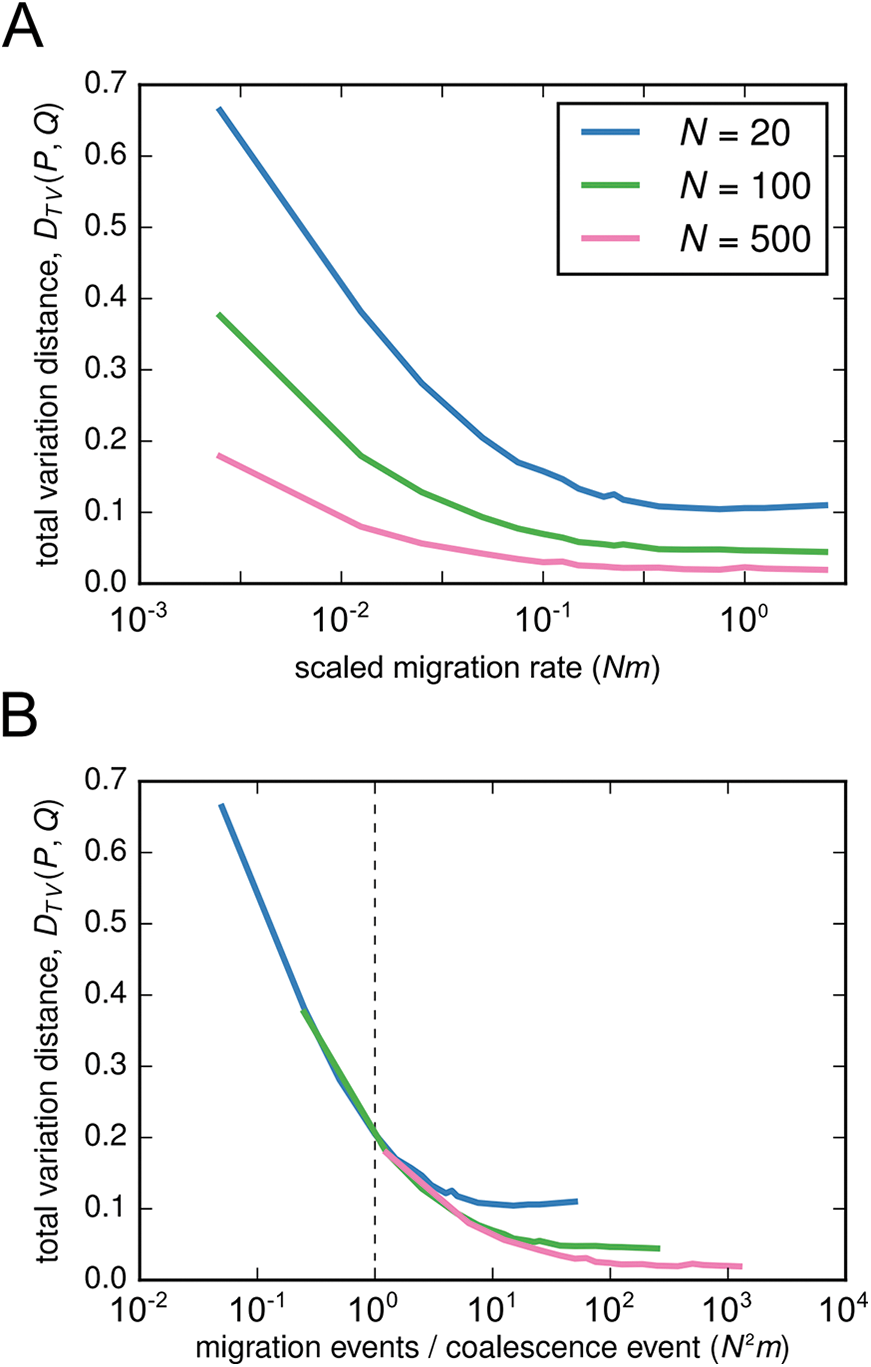
Mean total variation distance between distributions of coalescence times in two-deme population pedigrees and distributions of coalescence times predicted under a two-deme Wright-Fisher model. For each of a range of migration rates, twenty replicate pedigrees were simulated, and for each pedigree a distribution of coalescence times for a randomly sampled pair of individuals was calculated, and a total variation distance was calculated from the distribution that would be predicted under an analogous two-deme Wright-Fisher model. Each line shows how the mean of these total variation distances varies with migration rate. In Panel **A)**, total variation distance is shown against the average number of migration events per generation *Nm*. In Panel **B)**, total variation distance is shown against the mean number of migration events per coalescence event, *N* ^2^*m*. The dotted line shows where *N* ^2^*m* = 1.

### 2.3. Recent migrant ancestry and coalescence distributions in pedigrees

As is also the case for panmictic populations, the details of the *recent* sample pedigree are most important in determining the patterns of genetic variation in the sample. In panmictic populations, overlap in ancestry in the recent past increases the probability of coalescence in the very recent past, resulting in identity-by-descent. In structured populations, an individual may also have recent relatives from another deme. When this occurs, the distribution of pairwise coalescence times is potentially very different from the prediction in the absence of recent migrant ancestry due to the migration paths in the pedigree that lead to a recent change in demes. The degree of the difference in distributions is directly related to the amount of migrant ancestry, with more recent migration events causing
greater changes in the coalescence time distribution (Fig. 5).

**Figure 5:**
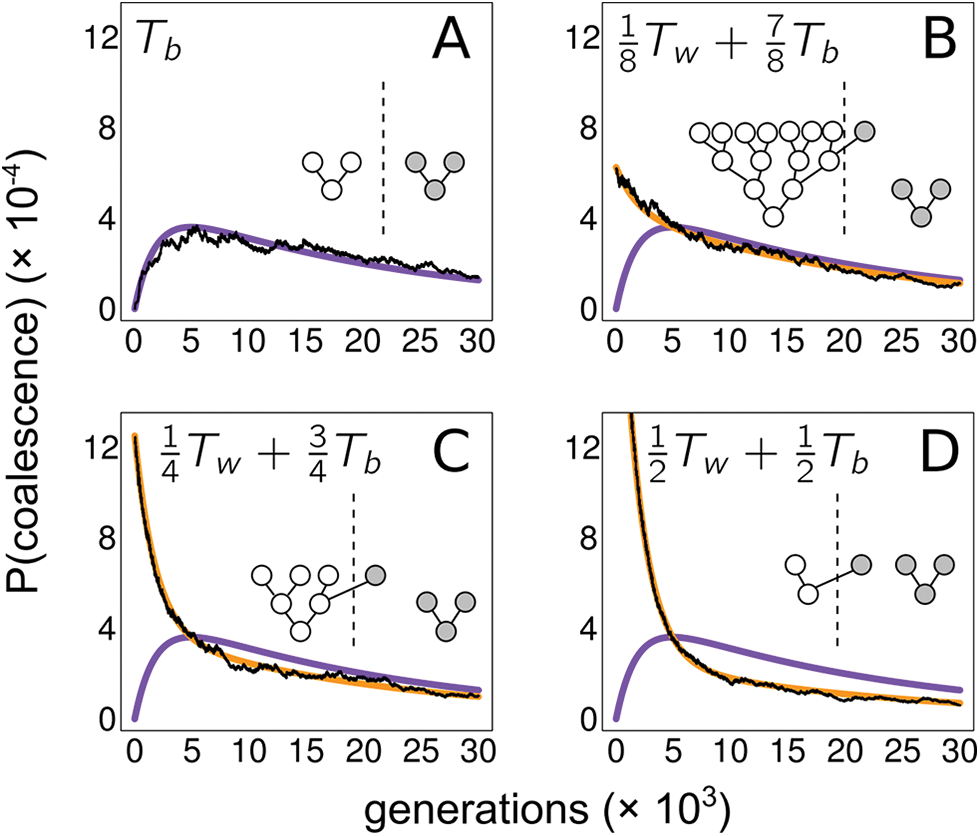
Pairwise coalescence time distributions for samples with migrant ancestry. Each panel shows the distribution of pairwise coalescence times for a pair of individuals with a different sample pedigree. In each simulated population pedigree, there are two demes of size *N* = 1000, and the scaled migration rate is 4*Nm* = 0.1. In each panel, the population pedigree was simulated conditional on the sample having the pedigree shown in the panel. The purple line shows the between-deme coalescence time distribution that would be expected in the absence of recent migrant ancestry, and the gold line shows the the mixture of the within- and between-deme coalescence time distributions that corresponds to the degree of migrant ancestry. Black lines are numerically calculated (i.e., exact) coalescence time distributions for the simulated example pedigrees.

If an individual in the sample has a migration event in its recent pedigree, there will be some probability that a gene copy sampled from this individual was inherited via the path through the pedigree including this recent migration event. If this is the case, it will appear as if that gene copy was actually sampled from a deme other than the one it was sampled from. In this way, the recent sample pedigree “reconfigures” the sample by changing the location of sampled gene copies, with the probabilities of the different sample reconfigurations depending on the number and timing of migration events in the recent pedigree.

If gene copies are sampled at many independently segregating loci from individuals having migrant ancestry, samples at some loci will be reconfigured by recent migration events, and others will not. In this scenario, the sample itself can be modeled as a probabilistic mixture of samples taken in different configurations, and probabilities of coalescence can be calculated by considering this mixture. For example, consider a sample of independently segregating loci taken from two individuals related by the pedigree shown in Fig. 5C, where one of two individuals sampled from different demes has a grandparent from the other deme. The distribution of pairwise coalescence times for loci sampled from this pair resembles the distribution of 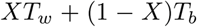, where *T_b_* is the standard between deme pairwise coalescence time for a structured-coalescent model with two demes, *T_w_* is the corresponding within deme pairwise coalescence time, and *X ∼* Bernoulli(1*/*4) (Fig. 5C). This particular mixture reflects the fact that a lineage sampled from the individual with migrant ancestry follows the path of migration through the pedigree with probability 1*/*4.

This sample reconfiguration framework can also be used to model identity-by-descent (IBD) caused by very recent coalescence due to branch overlap in the recent sample pedigree. If the pedigree causes an IBD event to occur with probability Pr(IBD), then the pairwise coalescence time is a mixture of the standard distribution (without IBD) and instantaneous coalescence (on the coalescent timescale) with probabilities 1 Pr(IBD) and Pr(IBD), respectively. If there is both recent coancestry and recent migration in the sample pedigree (or if there are multiple migration or coancestry events), the sample can be modeled as a mixture of several sample reconfigurations (e.g., Fig. 6).

**Figure 6:**
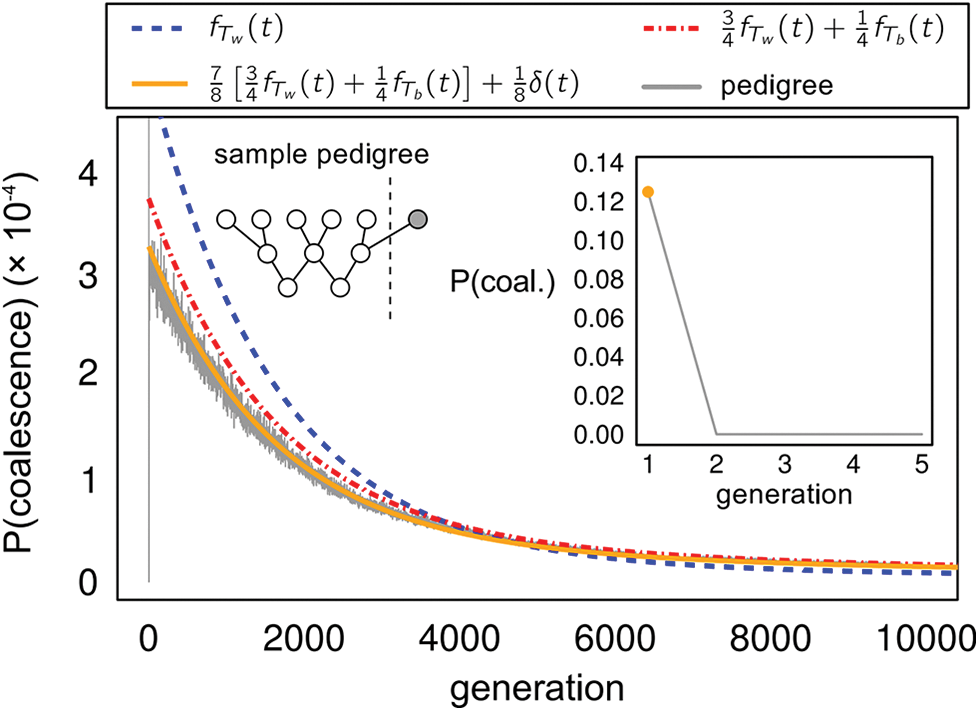
Distribution of pairwise coalescence times for a sample whose pedigree contains both recent migration and recent pedigree coancestry. The recent pedigree of the sample is shown, with the two sampled individuals located at the bottom of the pedigree. The distribution for the simulated pedigree (gray line) is calculated numerically from a pedigree of two demes of size *N* = 1000 each, with migration rate *M* = 4*Nm* = 0.2. The colored lines show mixtures of the within-deme coalescence time distribution (*f_T_w__*) and the between-deme distribution (*f_T__b_*). The inset shows the probability of coalescence during the first five generations; the probability mass at generation 1 (orange circle) is predicted by the mixture model accounting for both recent coancestry and migration (orange line). The red line shows the distribution if recent coancestry is ignored, and the blue line shows the distribution if both coancestry and migration are ignored.

This approach to modeling the sample implicitly assumes that there is some threshold generation separating the recent pedigree, which determines the mixture of sample reconfigurations, and the more ancient pedigree, where the standard coalescent models are assumed to hold well enough. The natural boundary between these two periods is around log_2_(*N*) generations, since any pedigree feature more ancient than that tends to be shared by most or all of the population (making such features “population demography”), whereas any more recent features are more likely to be particular to the sample.

We note that there is a long history in population genetics of modeling genetic variation in pedigrees as a mixture of different sample reconfigurations. Wright (1951) wrote the probability of observing a homozygous *A*_1_*A*_1_ genotype as 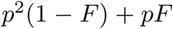, where *p* is the frequency of *A*_1_ in the population and *F* is essentially the probability of IBD calculated from the sample pedigree. This can be thought of as a probability for a mixture of two samples of size *n* = 2 (with probability 1 – *F*) and *n* = 1 (with probability *F*). The popular ancestry inference program STRUCTURE (Pritchard *et al.*, 2000) and related methods similarly write the likelihood of observed genotypes as a mixture over different possible subpopulation origins of the sampled alleles. Here, motivated by our simulations of coalescence in pedigrees, we explicitly extend this approach to modeling coalescence. In the next section, we propose an approach to inferring population parameters such as mutation and migration rates jointly with features of the sample pedigree such as recent coancestry and recent migrant ancestry. In the Discussion, we further consider the similarities and differences between our inference approach and existing methods.

### 2.4. Joint inference of the recent sample pedigree and population demography

The sample reconfiguration framework for modeling genetic variation in pedigrees, described above, can be used to calculate how recent migrant ancestry and overlap in the pedigree bias estimators of population-genetic parameters. In Appendix A, we calculate the bias of three estimators of the population-scaled mutation rate *θ* = 4*Nµ* in a panmictic population when the recent sample pedigree contains recently related or inbred individuals. In Appendix B, we calculate the bias of a moments-based estimator of *M* due to recent migration events in the sample pedigree. We use simulations to confirm these calculations (Figs. S3,S4). In both cases, if the recent sample pedigree is known, it is straightforward to correct these estimators to eliminate bias.

It is uncommon that the recent pedigree of the sample is known, however, and if it is assumed known, it is often estimated from the same data that is used to infer demographic parameters. Ideally, one would infer long-term demographic history jointly with sample-specific features of the recent pedigree. In this section, we develop a maximum-likelihood approach to inferring relationships in the recent sample pedigree and recent migrant ancestry jointly with long-term scaled mutation and migration rates. The method uses the approach proposed in the previous section: the sample pedigree defines some set of possible outcomes of Mendelian segregation in recent generations, and the resulting, reconfigured sample is modeled by the standard coalescent process.

We model a population with two demes each of size *N*, with each individual having probability *m* of migrating to the other deme in each generation. We rescale time by 2*N* so that the rate of coalescence within a deme is 1 and the rescaled rate of migration per lineage is *M/*2 = 2*Nm*. We assume that we have sampled two copies of each locus from each of *n*_1_ (diploid) individuals from deme 1 and *n*_2_ individuals from deme 2. We write the total diploid sample size as *n*_1_ + *n*_2_ = *n* so that the total number of sequences sampled at each locus is 2*n*. We index our sequences with 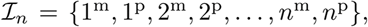 where *i*^m^ and *i*^p^ index the maternal and paternal sequences sampled from individual *i*. These are simply notational conventions; we do not assume that these maternal and paternal designations are known or observed.

The data 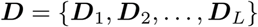 consist of genetic sequences sampled at *L* loci. We assume that there is free recombination between loci and no recombination within loci, and that mutation occurs according to the infinitely many sites model. We further assume that the ancestral versus derived status of each allele is known. Under these assumptions, the data at locus *i*, *D_i_*, can be stored as a 2*n × S_i_* binary matrix, where *S_i_* is the number of segregating sites at locus *i*. In this matrix, the first two rows pertain to the two sequences sampled from individual 1, the second two rows to the sequences sampled from individual 2, and so on. Since we assume that we do not know which sequences are maternal and which are paternal, the two rows pertaining to an individual are ordered arbitrarily in the matrix.

Each recent pedigree 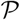 has some set of possible outcomes of segregation through the recent past, involving coalescence of lineages (due to recent pedigree relatedness or inbreeding) and movement of lineages between demes (due to recent migration events). The set of these sample reconfigurations is denoted 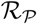, and each reconfiguration 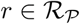 is a labeled partition of 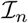, with the groups in each partition representing the lineages that survive after segregation back in time through the recent pedigree and the labels indicating the location (deme 1 or 2) of the lineage represented by the group. Corresponding to each sample reconfiguration 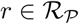, there is a probability 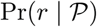 of that sample reconfiguration being the outcome of segregation back in time through the recent pedigree.

The joint likelihood of *θ*, *M*, and 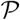 has the form

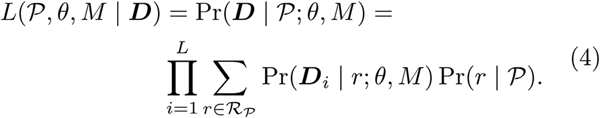

Probabilities are multiplied across independently segregating loci because their ancestries are assumed independent after conditioning on the recent sample pedigree. In order to calculate 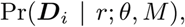 it is necessary to consider all the ways the sequences ***D**_i_* could have been inherited maternally and paternally, since we assume that this is unknown. For sequences 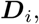 let 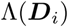 represent all 2*^n^* possible ways of labeling 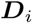 as maternal and paternal. Each such labeling associates each row (or sequence) in 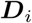 with an index in 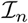. We then have

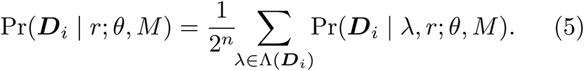

since each way of inheriting the sequences is equally likely to have occurred. Here we implicitly assume that the sequences for each individual are ordered (versus unordered) in 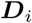. This has no effect on inference.

Since each reconfiguration 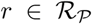 is a partition of *I_n_* and each labeling 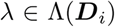 assigns each sequence a not assume that these maternal and paternal designations are known or observed. maternal-paternal label from 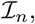 together the reconfiguration *r* and maternal-paternal labeling *λ* create a deme-labeled partition of the sequences 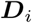, which we denote 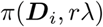. This partition represents the coalesced ancestral sequences after segregating back in time through the recent pedigree.

Each term in the sum in (5) corresponds to a sampling probability for one of these partitions. These are

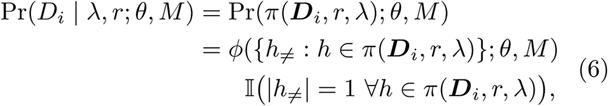

where 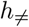 is the set of non-identical sequences in the collection of sequences 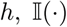 is an indicator function, and 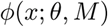 is the standard infinite-sites sampling probability of sequences *x*, calculated in the usual way without any reference to a pedigree. This sampling probability can be calculated numerically using a dynamic programming approach (Griffiths and Tavaré, 1994; Wu, 2010, see below).

The above equation says that conditional on certain sequences being IBD (*i.e.*, they are in the same group in the partitioned sequences), the sampling probability is the standard infinite-sites probability of the set of sequences with duplicate IBD sequences removed and the deme labelings of the different groups made to match any migration events that may have occurred. If any of the sequences designated as IBD are not in fact identical in sequence, the sampling probability for that reconfiguration and maternal-paternal labeling is zero. This is equivalent to assuming that no mutation occurs in the recent part of the pedigree.

Together, (4), (5), and (6) give the overall joint log-likelihood of mutation rate *θ*, migration rate *M*, and pedigree 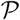 given sequences 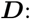

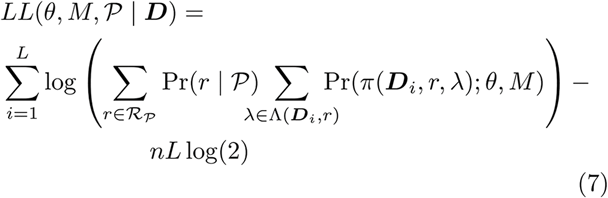

Our goal is to maximize (7) over *θ*, *M*, and 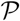 in order to estimate these parameters. The most direct approach would be to generate all possible recent sample pedigrees and maximize the log-likelihood conditional on each pedigree in turn. Logically, it would make sense to consider all sample pedigrees extending back to generation ∼ log_2_(*N*), since pedigree ancestry converges on this timescale (Chang, 1999; Rohde *et al.*, 2004) and this is when the pedigree affects probabilities of coalescence the most. However, with this approach the number of pedigrees to consider would be prohibitively large, and even enumerating all such pedigrees would be a difficult problem (e.g., Steel and Hein, 2006). Even if only the first few generations back in time are considered, the number of possible sample pedigrees will be large, and many sample pedigrees will contain extensive overlap or many migration events and thus would be unlikely to occur in nature. In most populations, it is more likely the case that each individual in the sample will has few overlaps or migration events in the few most recent generations, if any.

With this in mind, we limit the set of possible sample pedigrees to only those sample pedigrees containing two or fewer events, counting both overlap and migration events. We further limit the number of possible sample pedigrees by considering only the first four generations back in time, counting the present generation as the first generation. This is fewer generations than the *∼* log_2_(*N*) generations until the pedigree converges, but increasing the maximum generation would increase the number of possible pedigrees to the point of exceeding computational resources. However, since events in the pedigree have the greatest effect on estimates of demographic parameters when they occur recently (Figs. S3,S4), considering only the most recent generations addresses the greatest sources of bias in demographic parameters due to the sample pedigree.

Finally, since we assume that we do not know the parental origin of each sequence, we further reduce the number of possible sample pedigrees by including only pedigrees that are unique up to labeling of ancestors as maternal and paternal.

To generate all two-deme, four-generation sample pedigrees with two or fewer events, we first generate all possible two-generation pedigrees by considering all the ways that *n* individuals (where *n* is the diploid sample size) can have parents in the two different demes, and then from each of these two-generation pedigrees, three-generation pedigrees were produced by doing the same, and then the four-generation pedigrees were produced from the three generation pedigrees. The sample reconfiguration distribution of each four-generation pedigree was calculated by enumerating all possible ways of partitioning ancestral lineages into different ancestral chromosomes in each ancestral generation, and each such partition was counted as a state in a sparse matrix containing transition probabilities state in a sparse matrix containing transition probabilities (calculated by the rules of Mendelian segregation) from one generation to the next. The generation-by-generation transition matrices were multiplied together to obtain the final sample reconfiguration distribution.

Not every pedigree in the set we consider produces a unique distribution of sample reconfigurations, so in our inference method, the pedigree is not strictly identifiable. Thus, we place the pedigrees into groups that produce the same distribution of sample reconfigurations, and these are taken as the domain of pedigrees in inference.

To calculate the standard sampling probabilities needed in (6), we use the dynamic-programming method described by Wu (2010). In principle, it should be possible to calculate the log-likelihood of all pedigrees simultaneously, since any reconfiguration of the sample by IBD or migration must correspond to one of the ancestral configurations in the recursion solved by Wu’s (2010) method (see also Griffiths and Tavaré, 1994). For particular values of *θ* and *M*, after solving the ancestral recursion only once (and storing the sampling probabilities of all relevant ancestral configurations), the likelihood of any pedigree can be calculated by extracting the relevant probabilities from the recursion. However, in order to take this approach to maximize the log-likelihood, it is necessary to solve the recursion on a large grid of *θ* and *M*. In practice, we find that it is faster to maximize the log-likelihood separately for each pedigree, using standard derivative-free numerical optimization procedures to find the *M* and *θ* that maximize the log-likelihood for the pedigree.

To test our inference method we simulated datasets of samples from 1000 independently segregating loci, generated by simulating coalescence through a randomly generated pedigree of a two-deme population with deme size *N* = 1000 and one of two migration rates, *M ∈* {0.2, 2.0} We sampled one individual (two sequences at each locus) from each deme. Sequence data were generated using `coalseam` using one of two mutation rates: *θ/*2 = 0.5 (when *M* = 0.2) or *θ/*2 = 1.0 (when *M* = 2.0). In order to investigate the effects of recent migration events on the estimation on *θ* and *M*, each replicate dataset was conditioned on having one of three different sample pedigrees with differing amounts of recent migrant ancestry (see Fig. 7). We calculated maximum-likelihood estimates of *θ* and *M* for each of the 78 groups of pedigrees producing distinct reconfiguration distributions. We compared maximum-likelihood estimates to the estimates that would be obtained from a similar maximum-likelihood procedure that ignores the pedigree (*i.e.*, assuming the null pedigree of no sample reconfiguration).

**Figure 7:**
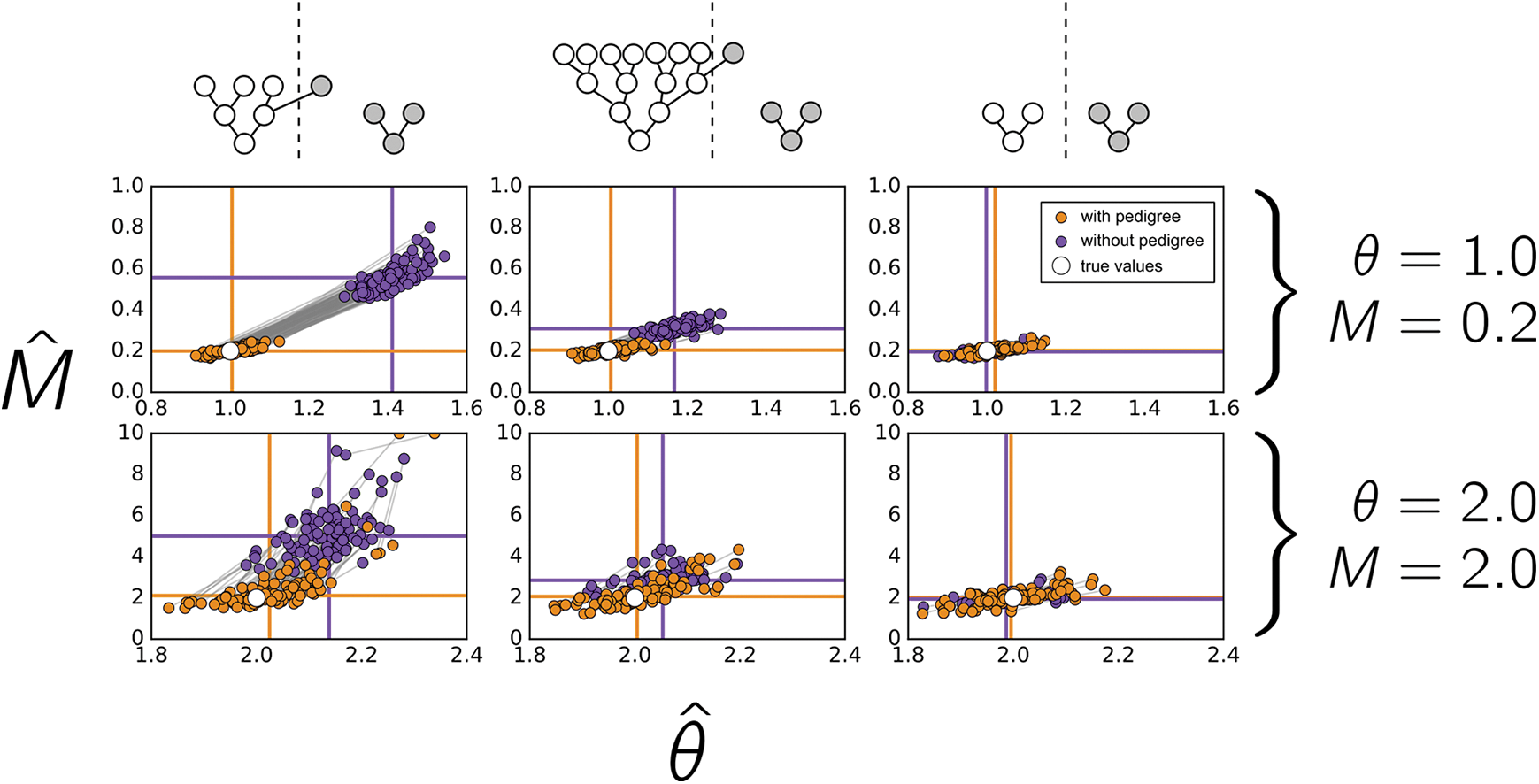
Maximum-likelihood mutation rates and migration rates for datasets simulated through different sample pedigrees. Each point depicts the maximum-likelihood estimates of *θ* and *M* for a particular simulation. Orange points show estimates obtained when the pedigree is included as a free parameter, and purple points show estimates obtained when the pedigree is assumed to have no effect on the data. In the first two columns, one of the sampled individuals is conditioned upon having a relative from the other deme, and in the third column the data are generated from completely random pedigrees. In each panel the true parameter values are shown with a solid white circle, and horizontal and vertical lines show means across replicates. Gray lines connect estimates calculated from the same dataset.

When there was recent migrant ancestry in the sample, assuming the null pedigree to be the true pedigree produced a bias towards overestimation of the migration rate (Fig. 7), since the early probability of migration via the migration path must be accommodated by an increase in the migration rate. For this reason, the overestimation of the migration rate was greater when the degree of recent migrant ancestry was greater. The population-scaled mutation rate was also overestimated when the recent migrant ancestry in the pedigree was ignored, presumably because migration via the migration path did not decrease allelic diversity as much as the overestimated migration rate should. Including the pedigree as a free parameter in the estimation corrected these biases. Estimates from simulations of samples lacking any features in the recent pedigree produced approximately unbiased estimates of *θ* and *M* (Fig. 7).

The pedigree was not inferred as reliably as the mutation and migration rates (Fig. 8). When the simulated pedigree contained a recent migration event, the estimated pedigree was the correct pedigree (out of 78 possible pedigrees) roughly 20–25% of the time. For pedigrees with no migrant ancestry and no relatedness, the correct pedigree was inferred about one third of the time. Most of the errors in pedigree estimation are due to mis-estimation of relatedness in the sample. If only the details of the migration ancestry are of interest, results are much better, with the correct migration ancestry inferred 50–100% of the time.

**Figure 8:**
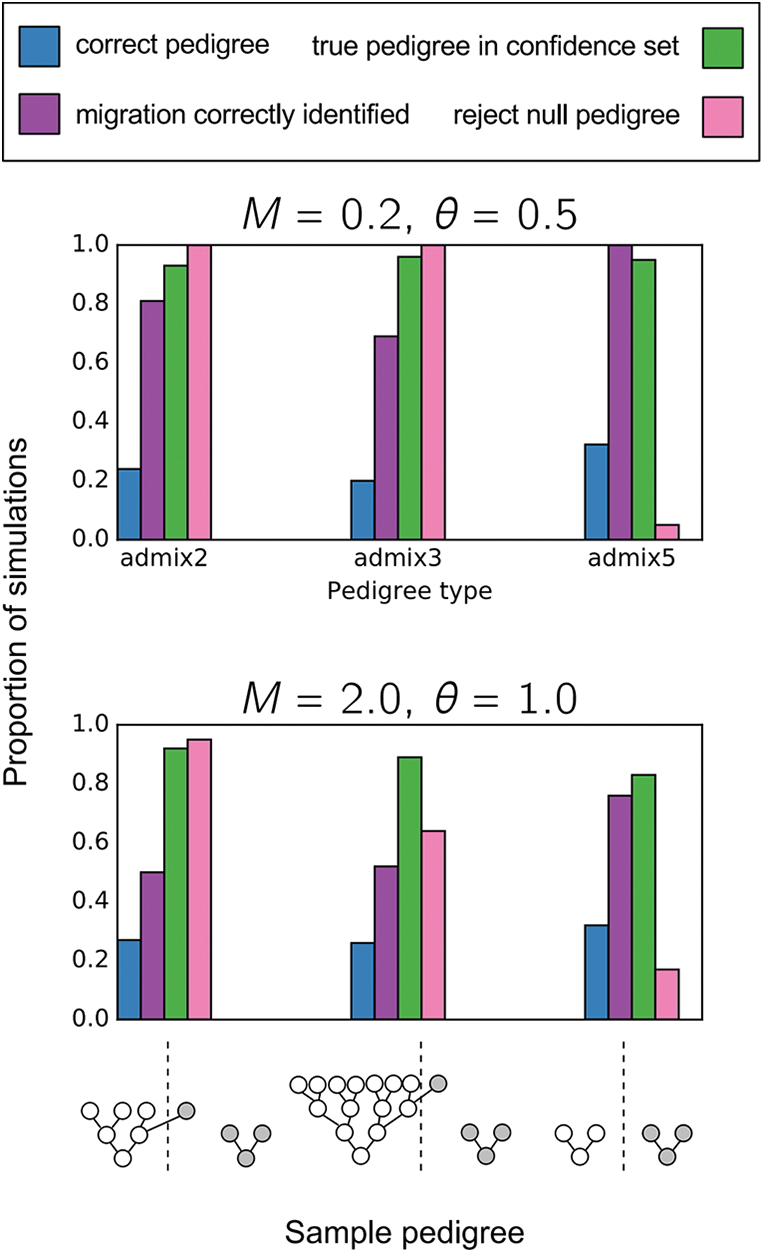
Inference of sample pedigrees. For simulations of 1000 infinite-sites loci, with ***θ*** = 0.5 and ***M*** = 0.2 (**A**) or ***θ*** = 1.0 and ***M*** = 2.0 (**B**), different measurements of the accuracy and power of pedigree inference are shown. The conditioned-upon sample pedigrees are shown at the bottom of the figure. Blue bars show the proportion of simulations in which the maximum-likelihood pedigree was the true pedigree. Purple bars show the proportion of simulations where it was inferred that sampled individuals had the correct amount of migrant ancestry. Green bars show the proportion of simulations in which the true pedigree was found within the approximate 95% confidence set of pedigrees, and pink bars show the proportion of simulations in which the null pedigree is rejected by a log-likelihood ratio test.

In addition to calculating a maximum-likelihood pedigree, it is possible to construct an approximate 95% confidence set of pedigrees using the fact that the maximum of the log-likelihood is approximately *χ*^2^ distributed when the number of loci is large. These pedigree confidence sets contained the true pedigree ∼ 83 − 96% of the time, depending on the true sample pedigree, mutation rate, and migration rate. A log-likelihood ratio test has nearly perfect power to reject the null pedigree for the simulations with the lesser migration rate; for simulations with the greater migration rate the power depended on the degree of migrant ancestry, with more recent migrant ancestry producing greater power to reject the null pedigree (Fig. 8). Type I error rates for simulations where the null pedigree is the true pedigree were close to *α* = 0.05.

Finally, we also simulated datasets of 1000 loci sampled from individuals with completely random pedigrees (i.e., sampled without conditioning on any migration events in the recent sample pedigree) in a small two-deme population of size *N* = 50 per deme with one of two different migration rates, 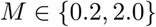. Whether or not the pedigree was included as a free parameter mostly had little effect on the estimates of *θ* and *M* (Fig. 9). However, in the few cases when the sample pedigree included recent migration events, the estimates were biased when the sampled pedigree was not considered as a free parameter, and including the sample pedigree in inference corrected this bias (Fig. 9).

**Figure 9:**
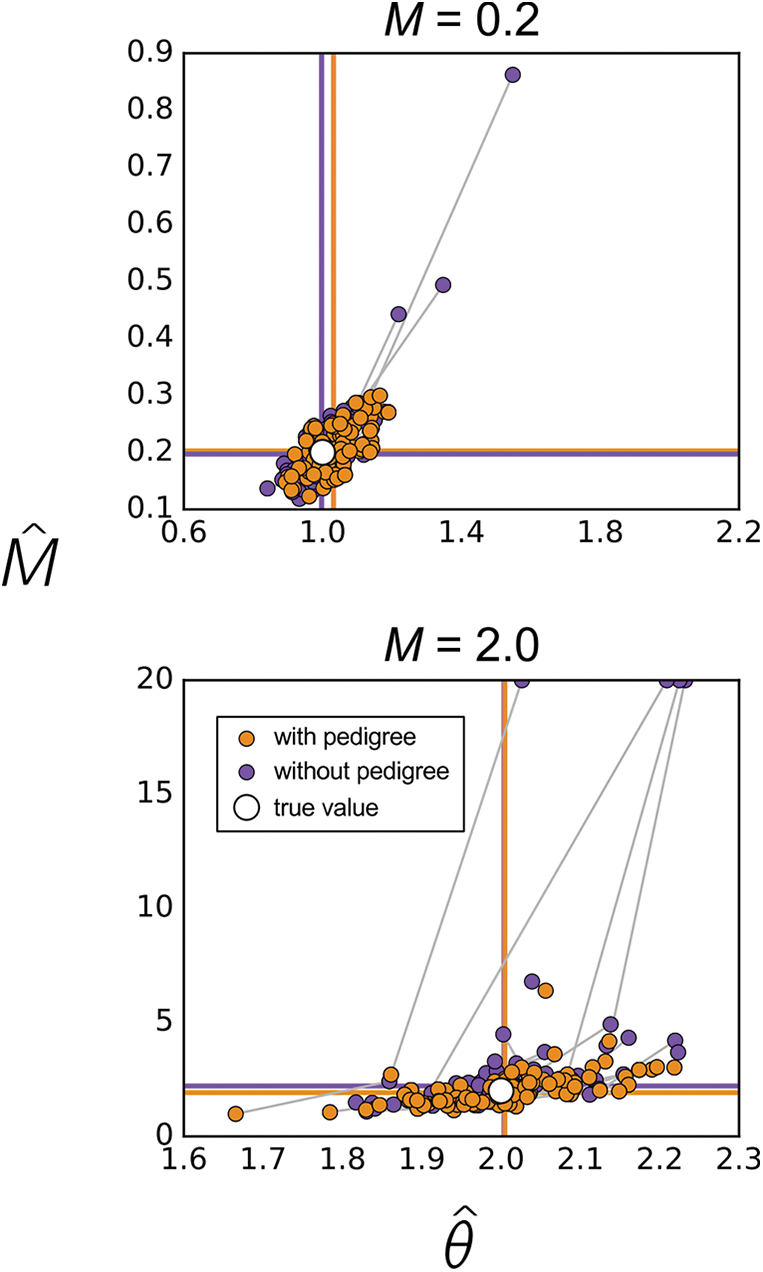
Maximum-likelihood mutation rates and migration rates for datasets of 1000 loci segregated through random pedigrees simulated in a two-deme population with deme size ***N*** = 50. Each point depicts the maximum-likelihood estimates of ***θ*** and ***M*** for a particular simulation. Orange points show estimates obtained when the pedigree is included as a free parameter, and purple points show estimates obtained when the pedigree is assumed to have no effect on the data. True parameter values are shown with white circles, and horizontal and vertical lines show means across replicates. Gray lines connect estimates calculated from the same dataset.

## 3. Discussion

Here we have explored the effects of fixed migration events in the population pedigree on the patterns of coalescence at independently segregating loci. In contrast to the case of panmictic populations, in structured populations the population pedigree can influence coalescence well beyond the time scale of ∼ log_2_(*N*) generations in the past. These effects are greater when the migration rate is small and are particularly pronounced when the total number of migration events occurring per generation is of the same order as the per-generation coalescence probability. When migration occurs more frequently than this (and there are no migration events in the recent sample pedigree), the particular history of migration events embedded in the population pedigree has less of an effect on coalescence, and the coalescence distributions based on the structured coalescent serve as good approximations to coalescent distributions within pedigrees.

We have also proposed a framework for inferring demographic parameters jointly with the recent pedigree of the sample. This framework considers how recent migration and shared ancestry events reconfigure the sample by moving lineages between demes and coalescing lineages. The inferred sample pedigree is the sample pedigree that has the maximum-likelihood mixture distribution of sample reconfigurations. In our implementation of this framework, we consider only sample pedigrees having two or fewer events, which must have occurred in the most recent four generations. At the expense of computational runtimes, the set of possible sample pedigrees could be expanded to include pedigrees with more events extending a greater number of generations into the past, but as these limits are extended, the number of pedigrees to consider grows rapidly. Another approach one could take would be to consider all of the reconfigurations (rather than sample pedigrees) with fewer than some maximum number of differences from the original sample configuration, and find the mixture of reconfigurations that maximizes the likelihood of the data, without any reference to the pedigree that creates the mixture. Such an approach may allow greater flexibility in modeling the effects of the recent pedigree, but searching the space of possible mixtures of reconfigurations — the *k*-simplex, if *k* is the number of possible reconfigurations — would likely be more challenging than maximizing over the finite set of sample pedigrees in the method we have implemented.

The sample reconfiguration inference framework is complementary to existing procedures for inferring recent admixture and relatedness in structured populations. The popular program STRUCTURE (Pritchard *et al.*, 2000) and related methods (Raj *et al.*, 2014; Alexander *et al.*, 2009; Tang *et al.*, 2005) are powerful and flexible tools for inferring admixture and population structure. Likewise, the inference tools RelateAdmix (Moltke and Al-brechtsen, 2014), REAP (Thornton *et al.*, 2012), and KING-robust (Manichaikul *et al.*, 2010) all offer solutions to the problem of inferring relatedness in the presence of population structure and admixture. Perhaps the most similar in scope is the method of Wilson and Ran-nala (2003), which uses inferred ancestry proportions to estimate migration rates in the most recent generations. For input, each of these methods take genotypes at polymorphic sites, often biallelic SNPs, that are assumed to segregate independently. Likelihoods are calculated from the probabilities of observing the observed genotypes under the rules of Hardy-Weinberg equilibrium. These methods are well suited for samples of a large number of SNP loci sampled from a large number of individuals. The inference procedure we have implemented, on the other hand, is capable of handling a sample of only a few individuals (*n ≈* 4), and likelihood calculations in our method are based on the coalescent in an explicit population genetic model, the parameters of which being the primary objects of inference. The pedigree is also explicitly modeled. While we have shown that our approach works well when its assumptions hold, if the narrow assumptions of our model do not hold, or if the primary goal of inference is to infer recent features of the sample pedigree *per se*, other, more flexible methods are likely to offer better performance.

Underlying our inference method is a hybrid approach to modeling the coalescent. Probabilities of coalescence are determined by the sample pedigree in the recent past, and then the standard coalescent is used to model the more distant past. This is similar to how Bhaskar *et al.* (2014) modeled coalescence when the sample size approaches the population size. In such a scenario, they suggest using a discrete-time Wright-Fisher model to model coalescence for the first few generations back in time and then use the standard coalescent model after the number of surviving ancestral lineages becomes much less than the population size. We note that this approach still implicitly marginalizes over pedigree relationships and that in a situation where the sample size nears the population size in a diploid population, there will be numerous common ancestor events and migration events in the recent sample pedigree. Thus in this context it may be important to consider the pedigree.

The sample reconfiguration framework could be extended to models that allow the demography of the population to vary over time (e.g., Gutenkunst *et al.*, 2009;

Kamm *et al.*, 2015). In such an application, if only a few individuals are sampled, it would be important to distinguish between the effects of very recent events that are more likely to be particular to the sample and the effects of events that are shared by all individuals in the population. The latter category of events are more naturally considered demographic history. On the other hand, if a sizable fraction of the population is sampled, inferred pedigree features may be used to learn more directly about the demography of the population in the last few generations. As sample sizes increase from the tens of thousands into the hundreds of thousands and millions (Stephens *et al.*, 2015), it will become more and more possible to reconstruct large (but sparse) pedigrees that are directly informative about recent demographic processes.

Unexpected close relatedness is frequently found in large genomic datasets (e.g. Gazal *et al.*, 2015; Pemberton *et al.*, 2010; Rosenberg, 2006). It is common practice to remove closely related individuals (and in some cases, individuals with admixed ancestry) from the sample prior to analysis, but this unnecessarily reduces the amount of information that is available for analysis. What is needed is a fully integrative method of making inferences from pedigrees and genetic variation, properly incorporating information about both the recent past contained in the sample pedigree and the more distant past that is the more typical domain of population genetic demographic inference. Here, by performing simulations of coalescence through pedigrees, we have justified a sample reconfiguration framework for modeling coalescence in pedigrees, and we have demonstrated how this can be incorporated into coalescent-based demographic inference in order to produce unbiased estimates of demographic parameters even when there is recent relatedness or admixed ancestry amongst the sampled individuals.

### 4. Acknowledgments

The authors thank Seungsoo Kim, Mark Martinez, Janet Song, and Katherine Xue for their their assistance during the early phases of this project. This manuscript was improved by the comments of Léandra King, Julia Palacios, Jerome Kelleher, and two anonymous reviewers. Computing resources were provided by Research Computing Group at Harvard University.

## 6. Appendix A: Pedigrees and biased estimators of *θ*

Estimates of the population-scaled mutation rate *θ* = 4*Nµ* will be downwardly biased if there are overlapping branches in the recent pedigree of the sample, since sequences will be identical with an artificially inflated probability, and this resembles coalescence prior to any mutation between the two identical sequences.

Suppose that we sample two copies of a DNA sequence from each of *n* diploid individuals from a panmictic population. As in the main text, we index these sequences with 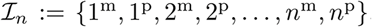 Let 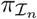 be the set of partitions of *I_n_*. Each pedigree 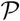 induces a set of sample reconfigurations 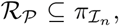 where each partition 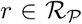 represents a possible outcome of segregation through the recent sample pedigree. Each reconfiguration 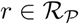 contains 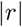 non-empty, disjoint subsets, each representing a distinct lineage that survives after segregating through the recent pedigree. Associated with each pedigree is also a probability distribution 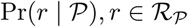 representing the Mendelian probabilities of the different sample reconfigurations.

We consider the bias of three estimators of *θ* that are unbiased in the absence of recent pedigree overlap. One estimator of *θ* we consider is Watterson’s (1975) estimator

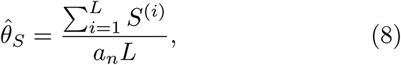

where *n* is the (haploid) sample size, *S*^(^*^i^*^)^ is the number of segregating sites at locus *i*, *L* is the number of sampled loci, and 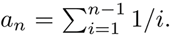 The expected value of 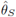 given pedigree 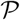 is

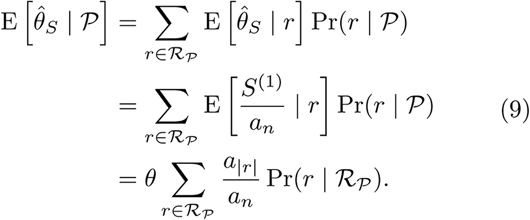

This follows from the fact that when there are *r* lineages surviving the recent pedigree, the expected number of segregating sites for that sample is 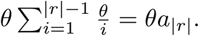

A second estimator of *θ* is 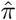, the mean number of differences between all pairs of sequences in a sample, which can be written in terms of the site-frequency spectrum:

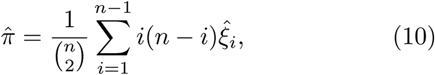

where 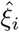 is the number of segregating sites present in *i* sequences in the sample.

To calculate the expected value of 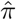 given 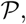 it is necessary to consider how overlap in the recent pedigree changes the site frequency spectrum. Define 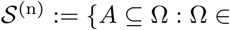 and 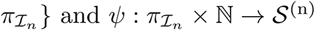 as

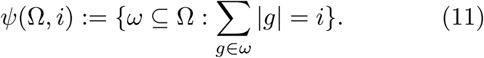

That is, 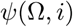 is the set of all subsets of the partition Ω such that the total size of all the groups in each subset is *i*. For example, for *n* = 3 and

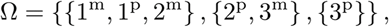

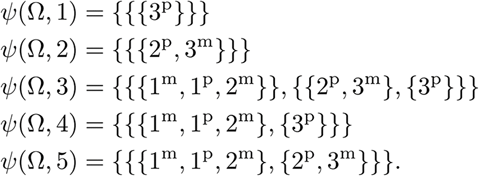

Then the expectation of 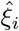 given reconfiguration 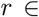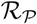 is

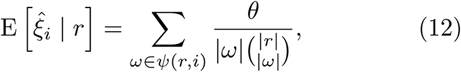

since each segregating site that is present in *i* present-day lineages must have occurred on the branch ancestral to the post-IBD lineages represented by some 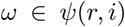. The expected number of mutations occurring on a branch subtending 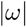 lineages is 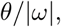 and the expected fraction of such mutations that occur on the branch ancestral to the lineages in 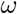 is 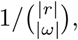 by exchangeability. This gives the expectation of 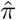 conditional on the pedigree:

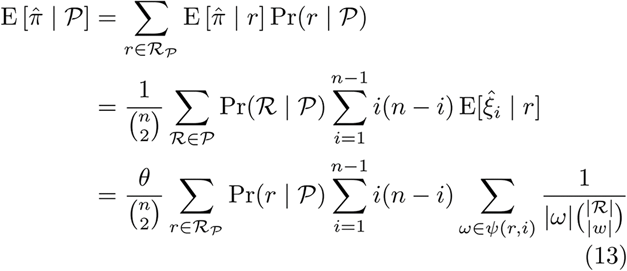

A third estimator of *θ* is 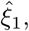 the number of singletons in the sample. The conditional expectation of 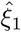 given a pedigree 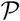 is

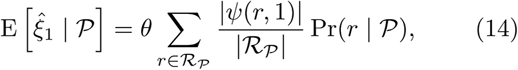

since only those mutations that occur on lineages that have not coalesced with any other lineages in the early pedigree can produce singletons.

To validate these calculations, we performed simulations of 200 loci sampled from individuals whose pedigree includes some amount of overlap. The simulations confirm the calculated biases for the different estimators of *θ* (Fig. S3).

## 7. Appendix B: Pedigrees and biased estimators of *M*

In a constant-sized structured population with two demes and a constant rate of migration 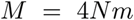 between demes, the expected within-deme and between-deme pairwise coalescence times are

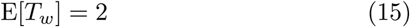

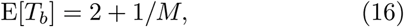

Let 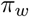 and 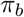 be the within-deme and between-deme mean pairwise diversity, respectively. Since 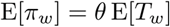 and 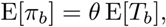 one estimator of *M* is

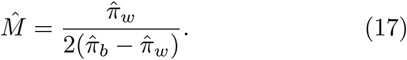

If some individuals in the sample have recent migrant ancestry, 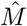 will be biased. In general it is not possible to calculate E[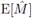], but it can be approximated by

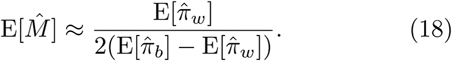

We sample two sequences from each of *n*_1_ individuals from deme 1 and *n*_2_ individuals from deme 2, defining 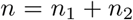 as the total (diploid) sample size. The sample is again indexed by 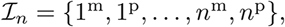 and we assume that the first 2*n*_1_ of these indices correspond to sequences sampled from deme 1 and the last 2*n*_2_ from deme 2.

In the context of a two-deme population, each group in the partitioned sample 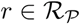 is labeled 1 or 2 to indicate which deme the lineage is found in after segregation back in time through the recent sample pedigree. For two-deme reconfiguration 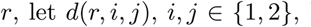 be a function that gives the number of lineages originally sampled from deme *i* that are found in deme *j* after segregation through the recent sample pedigree.

Assume that the recent sample pedigree contains migration events but no overlap. In this case, we can write the expectations of 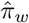 and 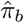 conditional on reconfiguration *r* as

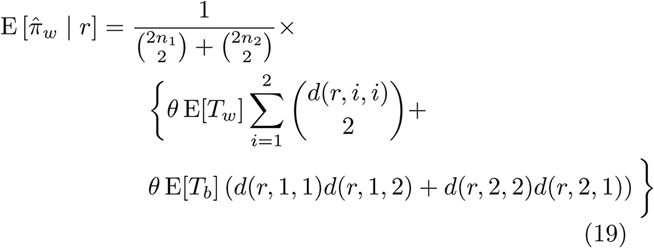

and

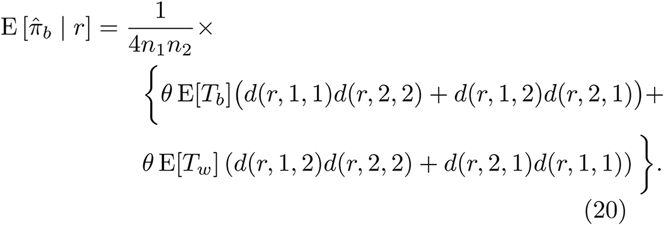

The approximate expectation of 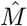 conditional on a pedigree 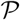 can be calculated using (19) and (20) together with

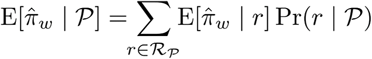

and

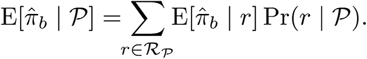

Simulations of infinite-sites loci taken from samples with a single migrant ancestor confirm these calculations (Fig. S4). This method of approximating 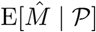 could be extended to accommodate recent sample pedigrees that contain both shared ancestry and migration, but we do not pursue this here.

## Supplementary Material

**Figure S1:**
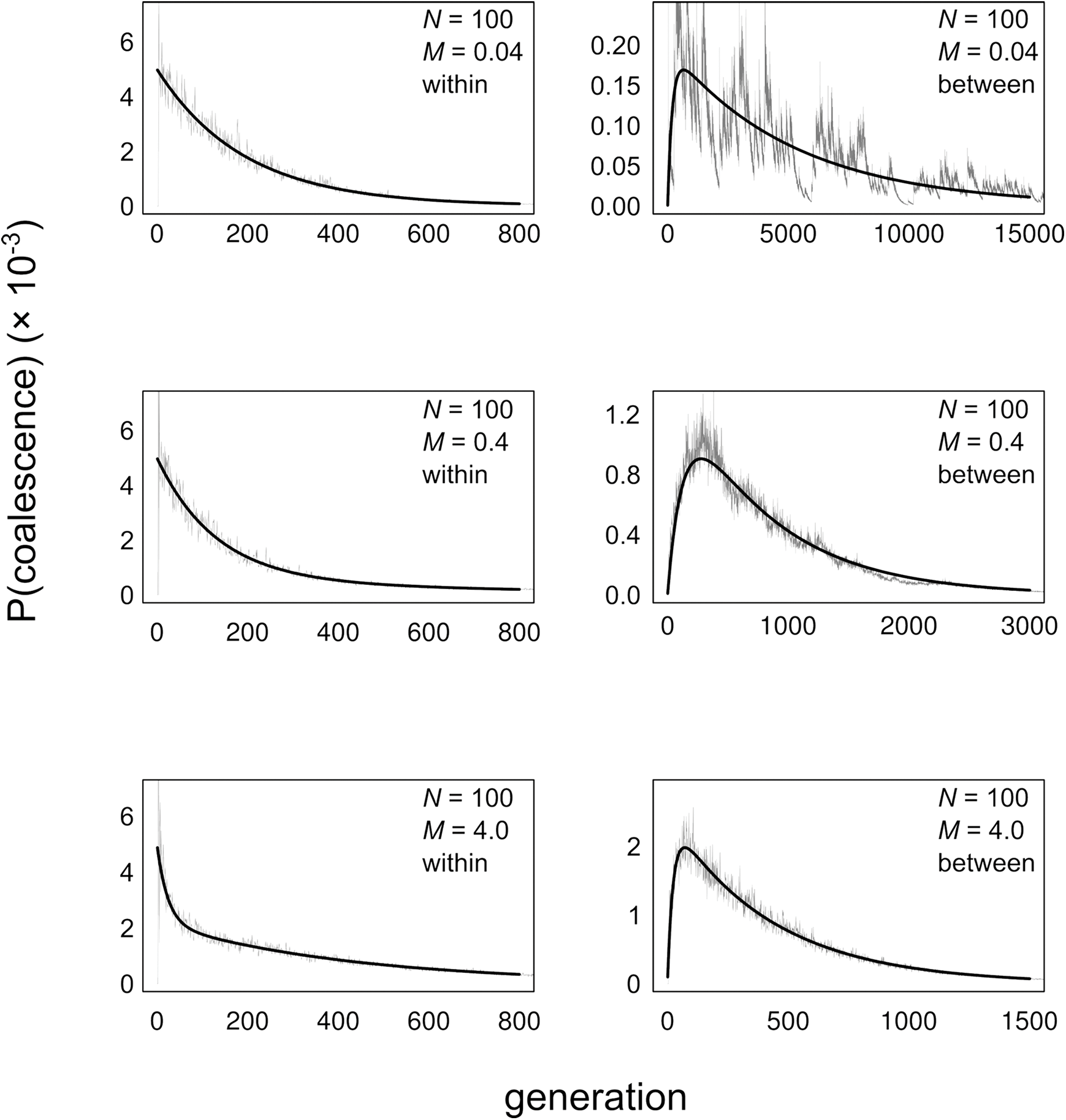
Distributions of coalescence times in two-deme populations with *N* = 100 individuals in each deme. Each panel shows a distribution of coalescence times for a particular value of the migration rate 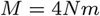 For panels in the left column, two individuals were sampled from the same deme (“within-deme” sampling), and in the right column two individuals were sampled from different demes.

**Figure S2:**
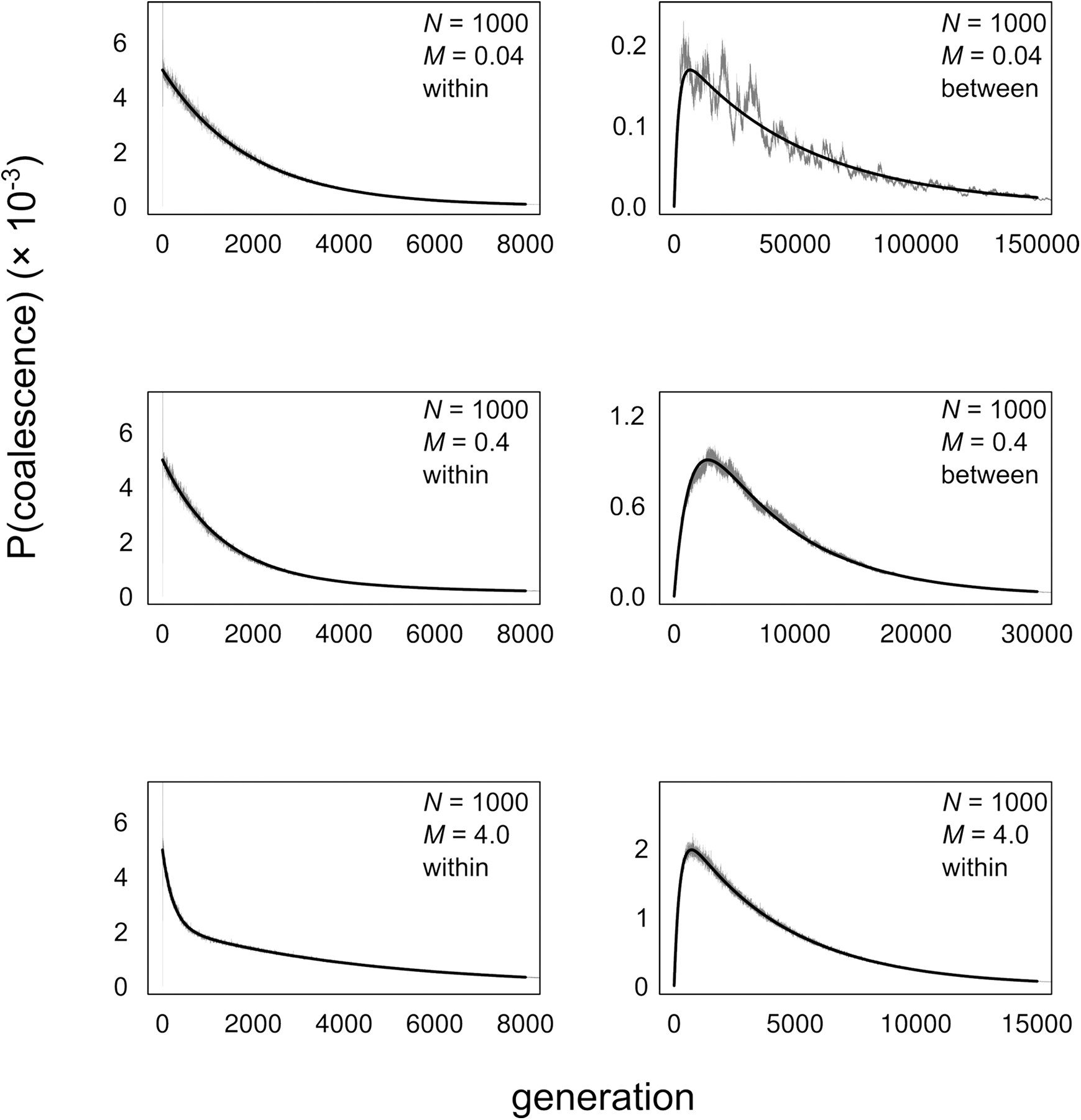
Distributions of coalescence times in two-deme populations with *N* = 1000 individuals in each deme. Each panel shows a distribution of coalescence times for a particular value of the migration rate 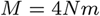. For panels in the left column, two individuals were sampled from the same deme (“within-deme” sampling), and in the right column two individuals were sampled from different demes.

**Figure S3:**
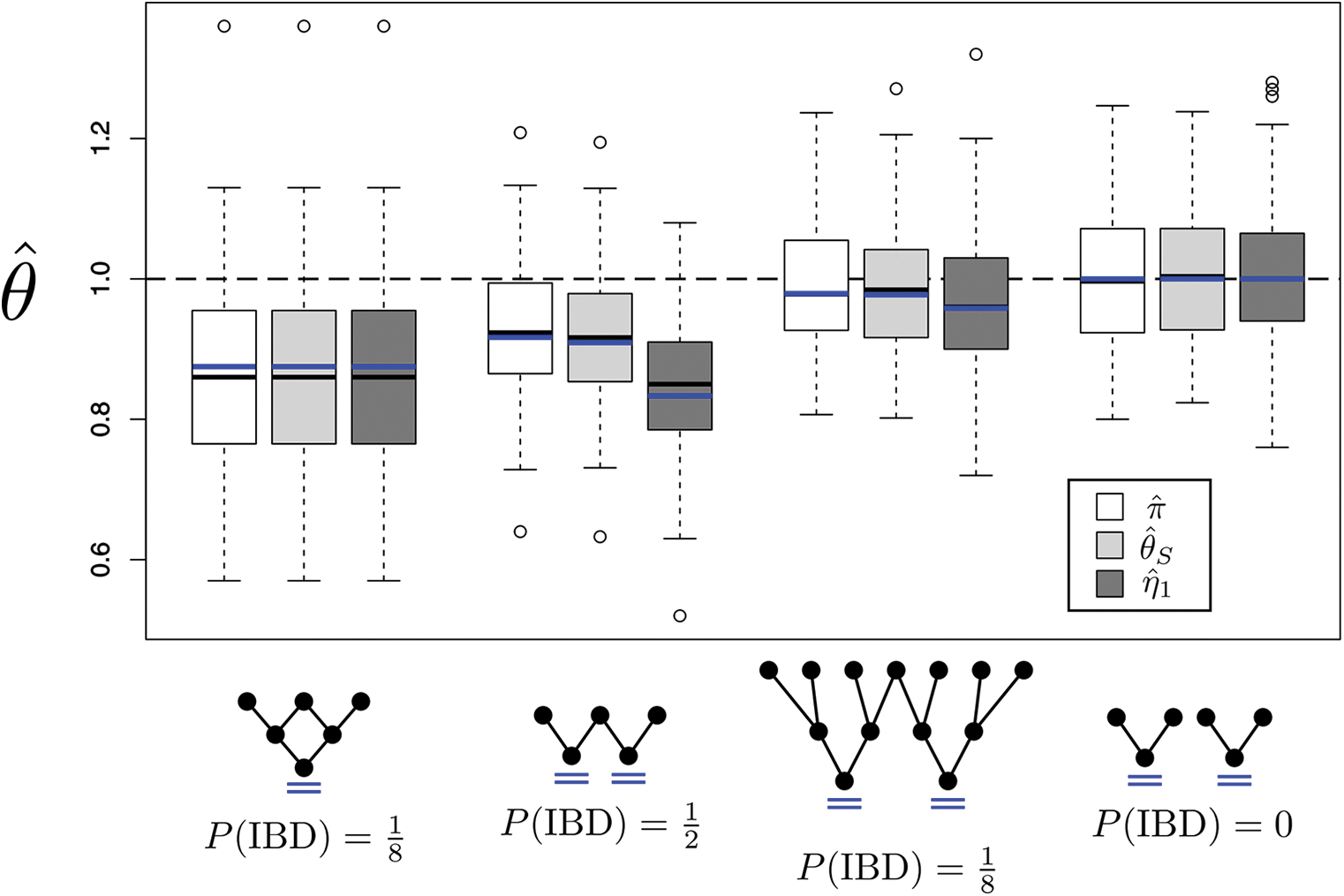
Estimates of 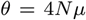 based on 100 replicate simulations of 200 loci segregating through sample pedigrees featuring identity by descent. Blue lines indicate theoretical predictions for individual pedigrees and estimators. Red horizontal lines indicate sample means across replicate simulations, and red vertical lines indicate twice the standard error of the mean. The true value of *θ* = 1.0 is indicated by the dashed line. Note that for the first pedigree, with *n* = 2, the three estimators are equivalent.

**Figure S4:**
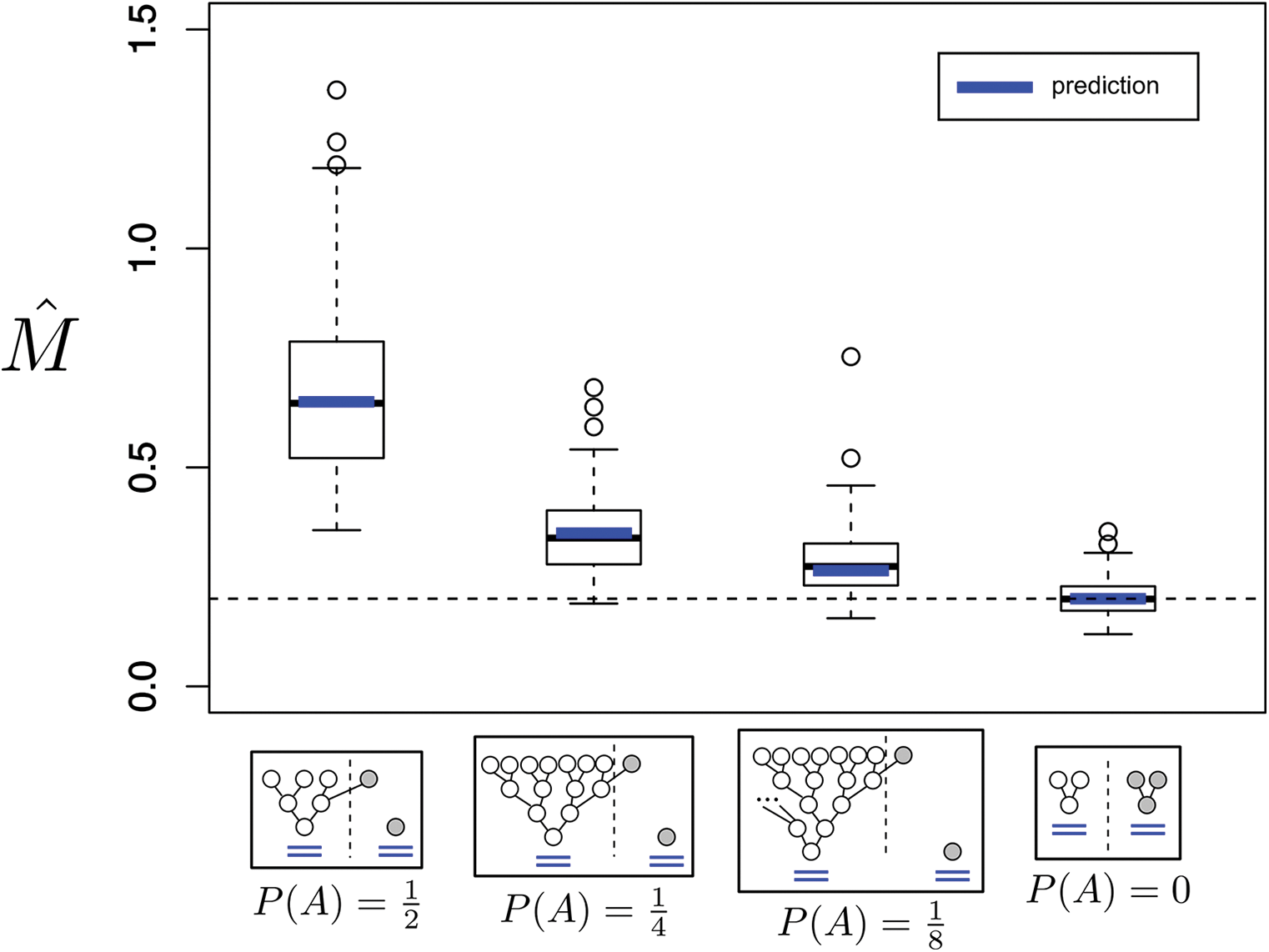
Estimates of *M* from the estimator given by (17) in replicate simulated genetic datasets of 100 loci segregating through sample pedigrees containing varying amounts of recent migrant ancestry. Blue lines indicate theoretical predictions for the different pedigrees. The true value of *M* = 0.2 is indicated by the dashed line. *P* (*M*) indicates the probability of migration in the indicated pedigrees.

